# Fully genetically encoded low-molecular-weight protein tags with defined shapes for direct molecular identification by cryo-electron tomography

**DOI:** 10.64898/2026.01.16.700029

**Authors:** Feng Luo, Rong Sun, Oliver Chalkley, Pingping Li, Qiangjun Zhou

## Abstract

Cryo-electron tomography (cryo-ET) enables three-dimensional visualization of cells in near-native states, but direct identification of specific proteins in situ remains challenging due to crowded cellular environments and the low intrinsic contrast of most proteins smaller than ∼500 kDa. Consequently, molecular identification often relies on indirect labeling strategies or bulky probes that can perturb native structures. Here we present a “shape-as-signal” strategy that uses fully genetically encoded protein tags with defined shapes as a molecular signal for direct identification by cryo-ET. We designed two single-chain, monomeric, low-molecular-weight tags: an extended V-shaped tag (62 kDa) and a compact triangular tag (85 kDa). Both adopt rigid geometries validated by cryo-electron microscopy and remain compatible with fluorescence microscopy when fused to fluorescent proteins. Their characteristic shapes are readily recognized and computationally detected in vitro. In cells, the V-shaped tag yields clear, non-disruptive signals at native locations. These results demonstrate that low-molecular-weight protein tags can be unambiguously detected and assigned in situ within crowded cellular environments. This single-step genetic tagging strategy enables seamless dual fluorescence and electron microscopy without exogenous probes, challenging the assumption that small protein tags are unsuitable for direct cryo-ET identification. More broadly, this approach establishes a scalable and minimally perturbative framework for visual proteomics and paves the way for multiplexed, shape-encoded molecular mapping in intact cells.

## Main

Light microscopy (LM) and electron microscopy (EM) reveal how proteins are organized and move in cells. Fluorescence microscopy (FM)—including modern super-resolution methods—can localize specific targets with 10-100 nm precision^1–7^, owing to protein and small-molecule fluorophores that enable selective labeling. EM provides a complementary view at higher spatial resolution. In particular, cryo-electron tomography (cryo-ET) visualizes cellular ultrastructure in three dimensions (3D) under near-native, vitrified conditions, resolving membranes, cytoskeletal elements, and large protein assemblies in intact cells^8–14^. However, cryo-ET is fundamentally limited by molecular identification. Only large, structurally distinctive complexes—such as ribosomes (>2.5 MDa), the 26S proteasome (∼2 MDa), mitochondrial respiratory supercomplexes (>1 MDa), and cytoskeletal polymers—are recognized directly in tomograms^15–20^. These complexes can be determined in situ at sub-nanometer resolution by subtomogram averaging (STA) only when they are abundant in cells. However, most proteins are <70 kDa and present at low abundance, making them difficult to identify effectively and precisely. As a result, cryo-ET often reveals where a structure resides in the cell, but not what it is. Precisely and unambiguously identifying these smaller, low-abundance proteins remains a central limitation of cryo-ET.

Multiple strategies have attempted to overcome this barrier by attaching high-contrast or physically large (>10 nm) markers to proteins of interest through affinity targeting or chemically induced coupling. Recent examples include nanogold particles^21^, iron-loaded ferritin cages^22,23^, DNA origami “signpost” scaffolds^24^, and multimeric protein tags such as genetically encoded multimeric particles (GEMs)^25^. These methods generate visible landmarks principally but accompanied with practical constraints: they often require post hoc labeling, have limited efficiency in cells, and can generate false positives through off-target binding, therefore limiting their general use. Cryogenic super-resolution fluorescence imaging improves localization of tagged proteins beyond the diffraction limit^26–34^, but correlation with cryo-ET is still only precise to tens of nanometers due to high fluorescence background and alignment error at cryogenic temperatures—typically not sufficient to assign identity to individual molecules^26,27,35,36^.

Here, we addressed this molecular identification limitation with a “shape-as-signal” strategy: developing a new class of genetically encoded, shape-defined protein tags that are directly visible by cryo-ET. Rather than attaching heavy metal particles or bulky scaffolds, we engineered low-molecular-weight, single-chain proteins that fold into rigid, geometrically distinctive 3D shapes intended to be recognizable by morphology (size and shape) alone. We engineered two single-chain, monomeric, shape-defined tags—a V-shaped protein (62 kDa) and a triangular protein (85 kDa)—whose rigid architectures were verified by single-particle cryo-EM. Using ferritin as a visibility benchmark in vitro, both tags produced clear densities in 3D cryo-tomograms, and STA resolved ferritin cages and tags. Inside the *Escherichia coli* (*E. coli)*, extended V-shaped protein architectures are inherently easier to distinguish around target assemblies, whereas more compact triangular designs, although still detectable, are more likely to be confused with surrounding punctate densities. In HeLa cells, fusion of both tags respectively to TOM70^NTD^ targeted them to the mitochondrial outer membrane without detectable trafficking or morphological defects; cryo-ET revealed a characteristic V-shaped density for enabling unambiguous, molecular-resolution (<2 nm) localization using standard 200 keV cryo-EM, while triangular tag signals were subtler but size-consistent. Fusion of GFP to the V-shaped tag provided dual optical and ultrastructural visibility, enabling broadly applicable correlative light and electron microscopy (CLEM) without the need for additional physical correlation steps. Together, these results establish a fully genetically encoded strategy for direct protein identification in cryo-ET and point toward a modular toolbox of shape-defined tags.

This strategy establishes a fundamentally new route for efficient, direct in situ protein labeling and opens opportunities to tackle an expanding set of scientific questions that demand both precise molecular mechanisms and intact ultrastructural context—critical areas that have long lacked practical solutions. It enables, for example, mapping the spatial arrangement of adaptor proteins or even individual protein isoforms at nascent membrane structures such as vesicles, distinguishing closely spaced paralogs within a membrane complex, and tracking assembly intermediates inside cells. Because the tags are fully genetically encoded, they can be seamlessly combined with standard molecular perturbations (mutants, truncations, rescue constructs), enabling coordinated functional and structural analyses and pushing visual proteomics toward single-molecule-level imaging in the native cellular context.

### Design and in vitro validation of V- and ***Δ***-shaped tags

To generate EM-visible protein tags with special geometries, we engineered an extended V-shaped tag (Fig. 1a-d) and a compact triangular tag (Fig. 1e-h).

**Fig. 1.**
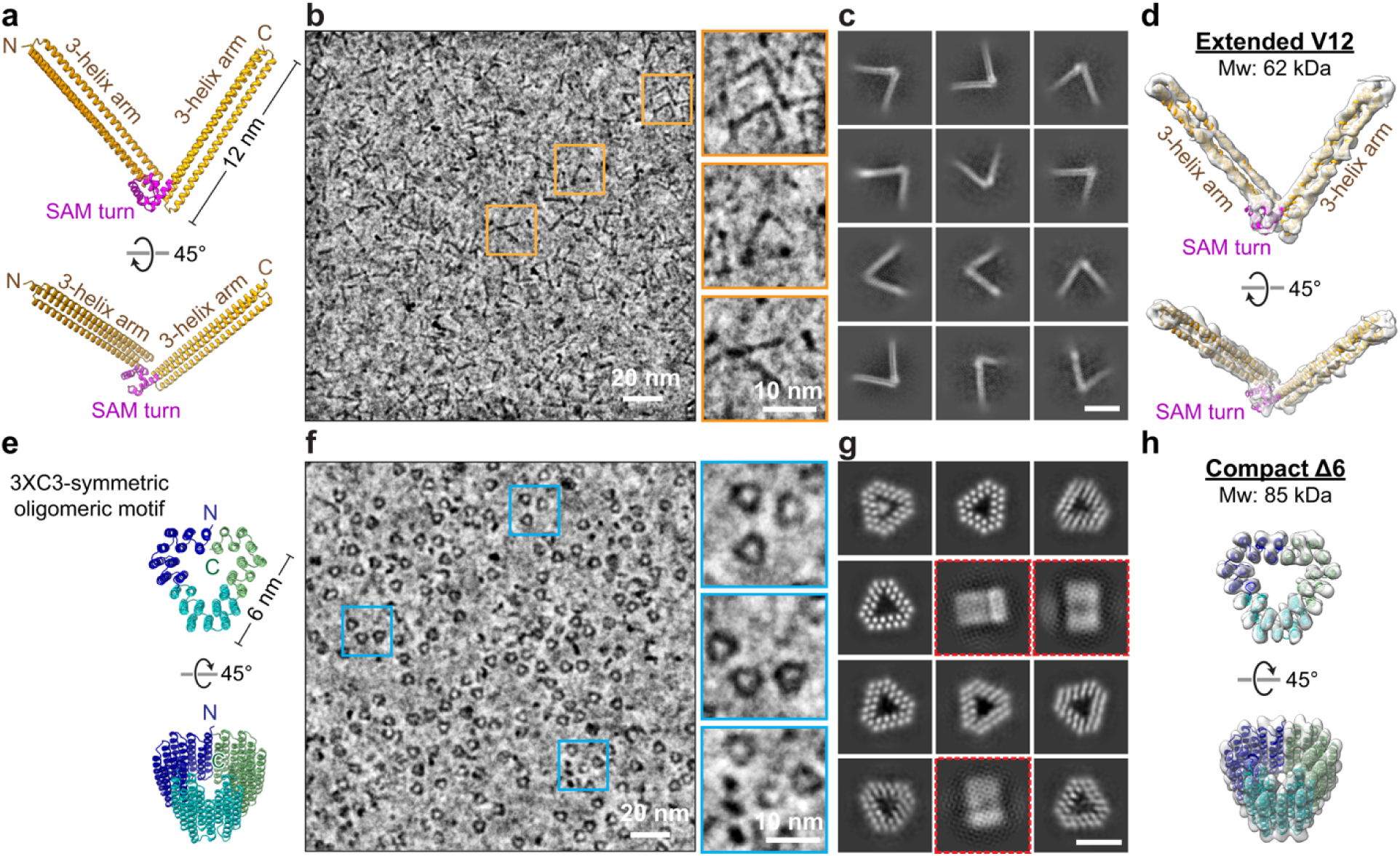
Design and structural characterization of V- and **Δ**-shaped protein tags. (a) Design of V-shaped tag (V12) predicted by AlphaFold2. Two three-helix arms connected by a rigid SAM-turn motif, forming an angle of ∼72° with an arm length of ∼12 nm. (b) Cryo-EM micrograph of purified V12 collected on a 200-keV Glacios cryo-TEM. Orange boxes mark representative V-shaped particles; enlarged views are shown at right. (c) Representative 2D class averages showing the characteristic V-shaped architecture, mostly in top view. Scale bar, 10 nm. (d) Cryo-EM density map of extended V12 (62 kDa) with the predicted model fitted into the density. (e) Design of the compact Δ6 tag predicted by AlphaFold2, consisting of three copies of C3-symmetric trimeric motif assembly ∼6 nm in diameter. (f) Cryo-EM micrograph of purified Δ6 collected on a 200-keV Glacios cryo-TEM. Blue boxes mark individual triangular particles; enlarged views are shown at right. (g) 2D class averages of Δ6 showing compact triangular geometries. Predominant top views are shown; side views are indicated by red boxes. Scale bar, 5 nm. (h) Cryo-EM density map and fitted predicted model of the compact Δ6 (85 kDa) reveal the expected triangular architecture.

For the V-shape, we built on a three-helix bundle scaffold^37^ and introduced rigid turn inspired by sterile α motif (SAM)^38^ to stabilize the angular junction between two bundles. Two bundles were connected by a rigid α-helical linker to form an extended V structure with a defined angle (Extended Data Fig. 1a). AlphaFold2^39,40^ predicted four V-shaped designs with ∼12 nm arms and inter-arm angles of approximately 60°, 72°, 90°, and 140° (Extended Data Fig. 1b). To preserve structural rigidity while preventing undesired interactions, we neutralized SAM oligomerization residues and turned surface hydrophilicity (Extended Data Fig. 2a). For the triangular tag, we adopted a C3-symmetric oligomeric motif^41^ as a structural template to generate an equilateral triangle with ∼6 nm sides (Fig. 1e). Surface residues were optimized for hydrophilicity to maintain solubility (Extended Data Fig. 2b). Thus, the V-shaped protein (∼62 kDa) forms extended 12 nm structures, whereas the triangular protein (∼85 kDa) adopts a compact ∼6 nm triangle (Figs. 1a, e and Extended Data Fig. 1b).

We expressed and purified all five designed constructs for single-particle cryo-EM analysis (SPA) to assess whether they folded as intended (Extended Data Fig. 3). Synthetic genes were cloned and expressed in *Escherichia coli (E. coli)*; proteins were purified by Ni-NTA immobilized metal affinity chromatography and analyzed by size-exclusion chromatography (SEC) to determine oligomeric state. Raw 200-keV cryo-EM micrographs showed well-defined particles for the 72° V-variant and the triangular construct; individual V- and triangular shapes were directly visible despite their low mass (Fig. 1b, f). As observed by EM and consistent with SEC, both proteins behaved as monomer with no detectable oligomerization (Fig. 1b, f). The other three V variants (60°, 90°, 140°) did not fold into the intended architectures. We designate the 12-nm 72° V-variant as V12 and the 6-nm triangular construct as Delta6 (Δ6). SPA reconstructions closely matched the designed models (Fig. 1c, d, g, h and Extended Data Fig. 4), confirming that both tags fold as intended and demonstrating our design strategy in which low-molecular-weight proteins are engineered to adopt defined geometries that enhance EM visibility. As is common for purified proteins, the samples exhibited preferred orientation on EM grids, with both V12 and Δ6 appearing predominantly in a “top” view (Fig. 1b, c, f, g).

### Tagging apoferritin cages in vitro

As a visibility benchmark, we fused V12 or Δ6 tag to *E. coli* ferritin (FtnA), which naturally assembles into a ∼12-nm nanocage composed of 24 subunits^42^. To minimize potential steric stress, we designed constructs containing two FtnA copies fused to either V12 (Extended Data Fig. 5a) or Δ6 (Extended Data Fig. 5c), such that a fully assembled cage could carry up to 12 tags. We expected peripheral tag densities surrounding the apoferritin cage both in vitro and in situ (Extended Data Fig. 5b, d). This design also enabled a direct test of whether tagging perturbs apoferritin cage assembly.

Tomograms of purified V12-ferritin revealed spherical cages with additional peripheral densities attributable to V12 (Fig. 2a). To further validate tagging, we evaluated several automated particle-picking pipelines^43–48^ and performed STA^46,47,49^. Notably, current cryo-ET picking and STA workflows are largely developed and optimized for large complexes (typically >500 kDa), which limits performance on low-molecular-weight features. Even so, using the deep-learning based program crYOLO^43,44^ and the template matching and correlation-based package PyTom^45,50^ (Extended Data Fig. 6a, b), particle picking followed by STA yielded independent averages for the apoferritin cage and V12 (Fig. 2b, c). The apoferritin cage was readily reconstructed to high resolution (5 Å), consistent with its size (∼12-nm outer diameter; ∼8-nm cavity; ∼465 kDa) and high symmetry (octahedral symmetry, 432 point group). By contrast, only 16.3% of V12 picks contributed to a low-resolution average (Fig. 2c, d), underscoring a known limitation: existing cryo-ET picking/STA pipelines, tuned for larger assemblies, struggle with low-molecular-weight targets like V12 due to low signal to noise ratio (SNR) and orientation ambiguity, as well as the inherent missing-wedge in cryo-ET^43–45,49,51^.

**Fig. 2.**
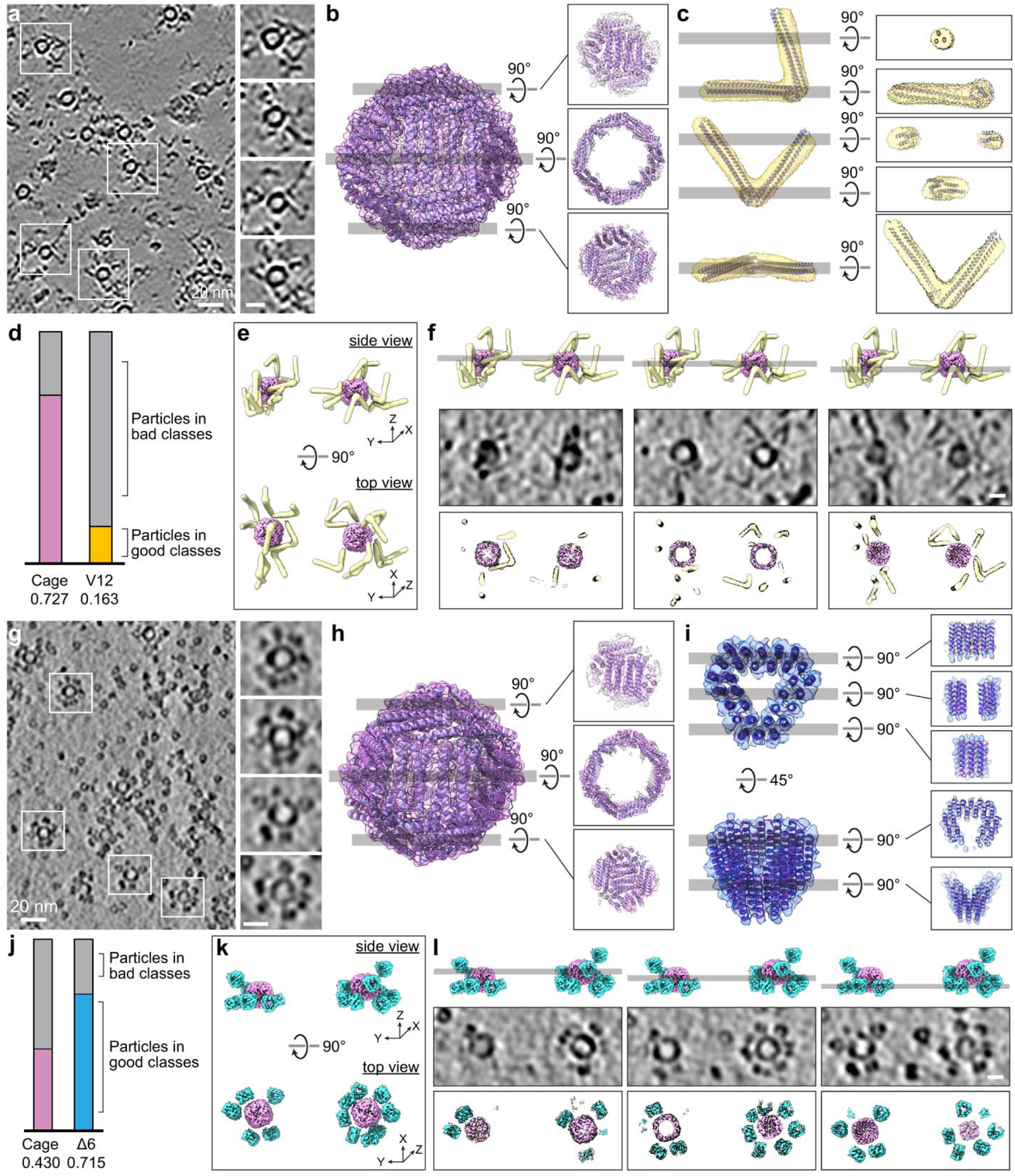
In vitro visualization and analysis of V12- and **Δ**6-tagged ferritin nanocages. (a) Representative cryo-electron tomogram slice of purified V12-tagged ferritin nanocages. Insets show enlarged regions highlighting individual cages and associated V-shaped densities. Scale bar, 20 nm (left) and 10 nm (right). (b, c) STA structures and representative orientated slice views of the ferritin cage (b) and the V12 tag (c), each reconstructed independently from purified tomograms. (d) Fraction of particles retained after classification for V12 (orange) and ferritin cages (magenta), illustrating the challenge of identifying small V-shaped tags in crowded tomograms. (e) Model of the ferritin cage and V12 tag obtained by STA and fitted into the 3D tomographic density. (f) Comparison of model and tomogram slices. Top, representative model slice corresponding to the tomogram slice; middle, tomogram slice; bottom, fitted model slices showing close agreement between model and density. Scale bar, 10 nm. (g) Representative cryo-electron tomogram slice of purified Δ6-tagged ferritin nanocages. Insets show enlarged regions highlighting individual cage and associated compact, triangular densities surrounding the cages corresponding to the Δ6 tag. Scale bar, 20 nm (left) and 10 nm (right). (h, i) STA structures and representative orientated slice views of the ferritin cage (h) and the Δ6 tags (i) reconstructed independently from purified tomograms. (j) Fractions of particles retained after classification for Δ6 (blue) and ferritin cages (magenta) showing that the compact triangular tags are more readily identified in vitro but may influence the structural analysis of target protein. (k) Model of the ferritin cage and Δ6 tag obtained by STA and fitted into the 3D tomographic density. (l) Comparison of model and tomogram slices. Top, representative model slice corresponding to the tomogram slice; middle, tomogram slice; bottom, fitted model slices showing close agreement between model and density. Scale bar, 10 nm.

In 2D tomographic slices, only views aligned near the V apex display a clear V (Fig. 2a, c); whereas most other orientations appear as two dots or a short line (Fig. 2a, c). In 3D, however, the V shape is evident: the averaged apoferritin cage and V12 volumes fit unambiguously into the tomographic densities, producing a coherent structural model (Fig. 2e and Supplementary Video 1), that confirms intact cage assembly and direct detectability of V12. Slice-wise densities agree with 2D projections of the fitted model (Fig. 2e, f), demonstrating that nearly the entire tags are visualized across orientations—further clarifying why existing particle picking and STA algorithms struggle with V12 despite its clear visibility in tomograms.

For Δ6-ferritin, tomograms likewise showed peripheral tag densities (Fig. 2g). Automated picking (crYOLO, PyTom; Extended Data Fig. 6c, d) and STA yielded independent averages for the cage and Δ6 (Fig. 2h, l), with a usable-particle fraction of 71.5% for Δ6 and 43.0% for the cage (Fig. 2j), underscoring the compact tag’s strong in vitro performance. Relative to V12-ferritin, the lower cage fraction of the cage in Δ6–ferritin datasets suggests that the compact Δ6 density may influence apoferritin cage picking. The averages recapitulated the expected geometries and fit perfectly into tomographic densities (Fig. 2k, l and Supplementary Video 2).

Slice views revealed triangular densities in top views and one or two discrete spots in side views, consistent with Δ6 orientation (Fig. 2i, l).

### Tagging apoferritin cages in E. coli

We next examined whether the V12 and Δ6 tags were detectable in situ. V12- and Δ6-tagged ferritin were expressed in *E. coli*, and 80–250-nm thick lamellae were prepared by a cryogenic focused ion-beam scanning electron microscope (cryo-FIB-SEM) (Extended Data Fig. 7a, b).

For V12-tagged ferritin, tomograms reconstructed with missing-wedge compensation and denoising using IsoNet^52^ revealed membranes, ribosomes, and numerous ∼12-nm nanocages (Fig. 3a, b). Template-based particle picking using PyTom (Extended Data Fig. 7c) followed by STA identified apoferritin cages in situ (Fig. 3d). Only a small fraction of particles contributed to the final average (Fig. 3e), underscoring the difficulty of detecting small features in crowded tomograms. Notably, close inspection of individual cages revealed extended densities consistent with the expected V-shaped geometry despite the tag’s modest mass (62 kDa). As anticipated, existing algorithms did not reliably detect or reconstruct the low-molecular-weight V12 tag in this context. Nevertheless, manual inspection consistently revealed V-shaped densities adjacent to nanocages—matching the in vitro structures and demonstrating direct recognition of V12 in situ (Fig. 3c, f, g and Supplementary Video 3).

**Fig. 3.**
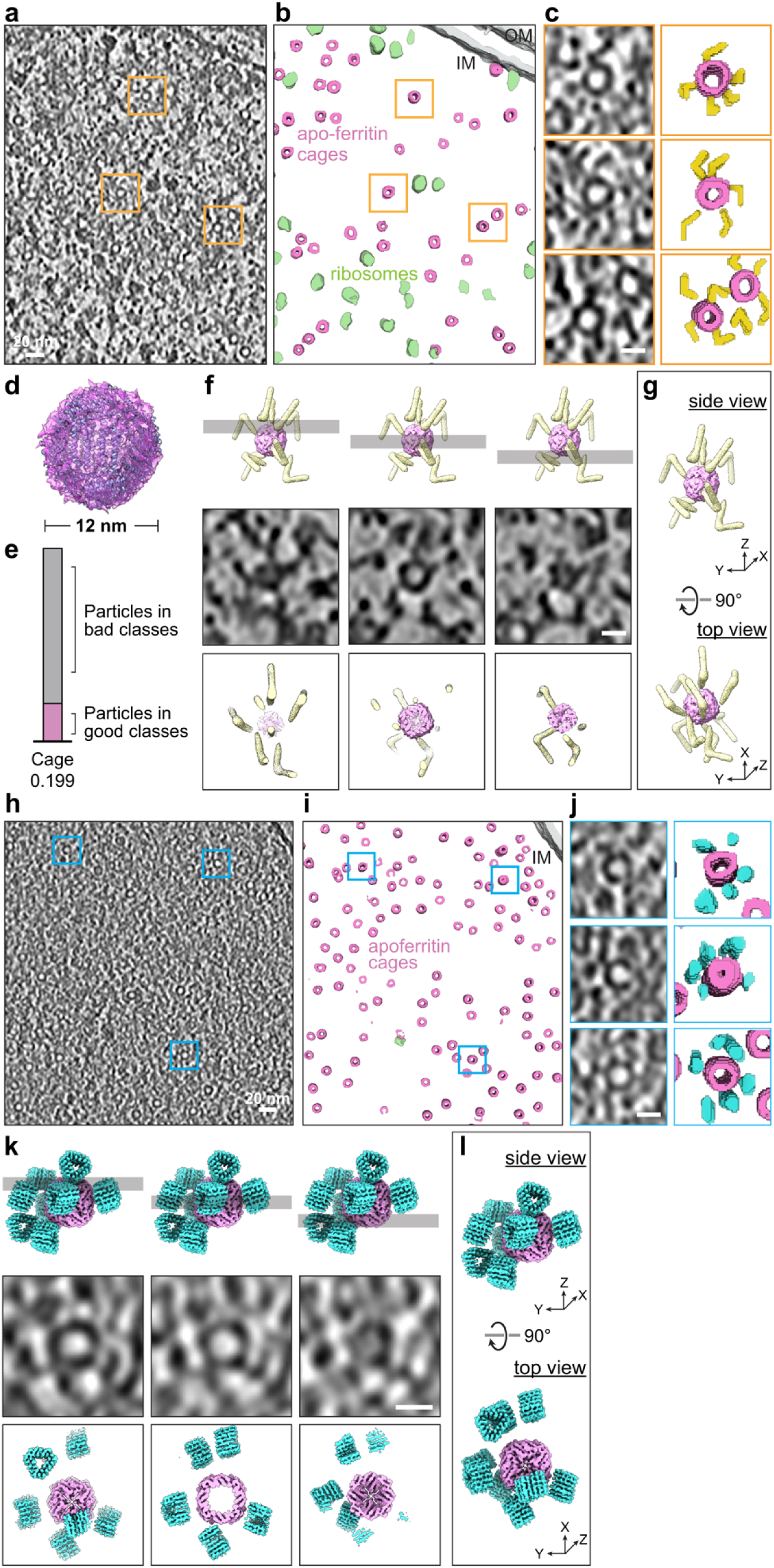
In situ visualization of V12- and **Δ**6-tagged ferritin cages in *E. coli* by cryo-ET. (a) Representative cryo-tomographic slice of a FIB-milled *E. coli* cell expressing V12-tagged ferritin nanocages. Orange boxes mark examples of nanocages. Scale bar, 20 nm. (b) Automated segmentation with Amira showing apo-ferritin cages (pink), ribosomes (green) within the cytoplasm and cell membranes (grey). (c) Enlarged views of boxed regions in panel (a) showing peripheral extended densities corresponding to V12 tags with annotated views at right. Scale bar, 10nm. (d) STA structure of the apo-ferritin cage from in situ particle picking. (e) Fraction of ferritin cage particles retained after in situ classification, illustrating the low yield of usable particles in crowded cellular environments. (f) Comparison of model and in situ tomogram slices. Top, representative model slice corresponding to the tomogram slice; middle, tomogram slice; bottom, fitted model slices showing close agreement between model and density. Scale bar, 10 nm. (g) Model of the ferritin cage and V12 tag obtained by in vitro STA and fitted into the in situ 3D tomographic density. (h) Representative cryo-tomographic slice of a FIB-milled *E. coli* cell expressing Δ6-tagged ferritin nanocages. Blue boxes mark examples of nanocages. (i) Segmentation highlighting apo-ferritin cages (pink). (j) Enlarged views of boxed regions in (h) showing compact peripheral densities corresponding to Δ6 tags with the annotation at right. Scale bar, 10nm. (k) Comparison of model and in situ tomogram slices. Top, representative model slice corresponding to the tomogram slice; middle, tomogram slice; bottom, fitted model slices showing close agreement between model and density. Scale bar, 10 nm. (l) Model of the ferritin cage and Δ6 tag obtained by in vitro STA and fitted into the in situ 3D tomographic density.

For Δ6-tagged ferritin, high quality of tomograms of cryo-FIB-milled *E. coli* likewise revealed nanocages (Fig. 3h, i). Around the cages, ∼5–6 nm dot-like densities were frequently observed (Fig. 3j-l and Supplementary Video 4), consistent with the compact triangular geometry of Δ6 and matching the in vitro structures (Fig. 2g-l). However, because similar punctate features are abundant throughout the cytoplasm, individual Δ6 tags, while detectable, were more prone to misidentification with surrounding densities.

Together, these results indicate that both tags could be detected in the crowded bacterial cytoplasm, with the extended V12 tag providing a more distinctive and recognizable shape cue than the compact Δ6 tag.

### Display on the mitochondrial surface in HeLa cells

Having confirmed the visibility of both tags in bacteria, we next tested their labeling performance on the mitochondrial surface in mammalian HeLa cells. To target a native membrane, V12 or Δ6 was fused to the N-terminal targeting fragment of (TOM70^NTD^)^25,53^ and appended GFP for fluorescence readout (Fig. 4a, b). Western blotting with anti-GFP confirmed robust expression of tagged constructs (Fig. 4c), and GFP fluorescence colocalized with the mitochondrial marker Hsp60 (Fig. 4d, m). Consistent results from anti-HA antibody and Mito-Tracker Red staining in HeLa cells, together with western blotting in HEK293T cells, confirmed proper expression and mitochondrial localization for both tags without detectable interference (Extended Data Fig. 8).

**Fig. 4.**
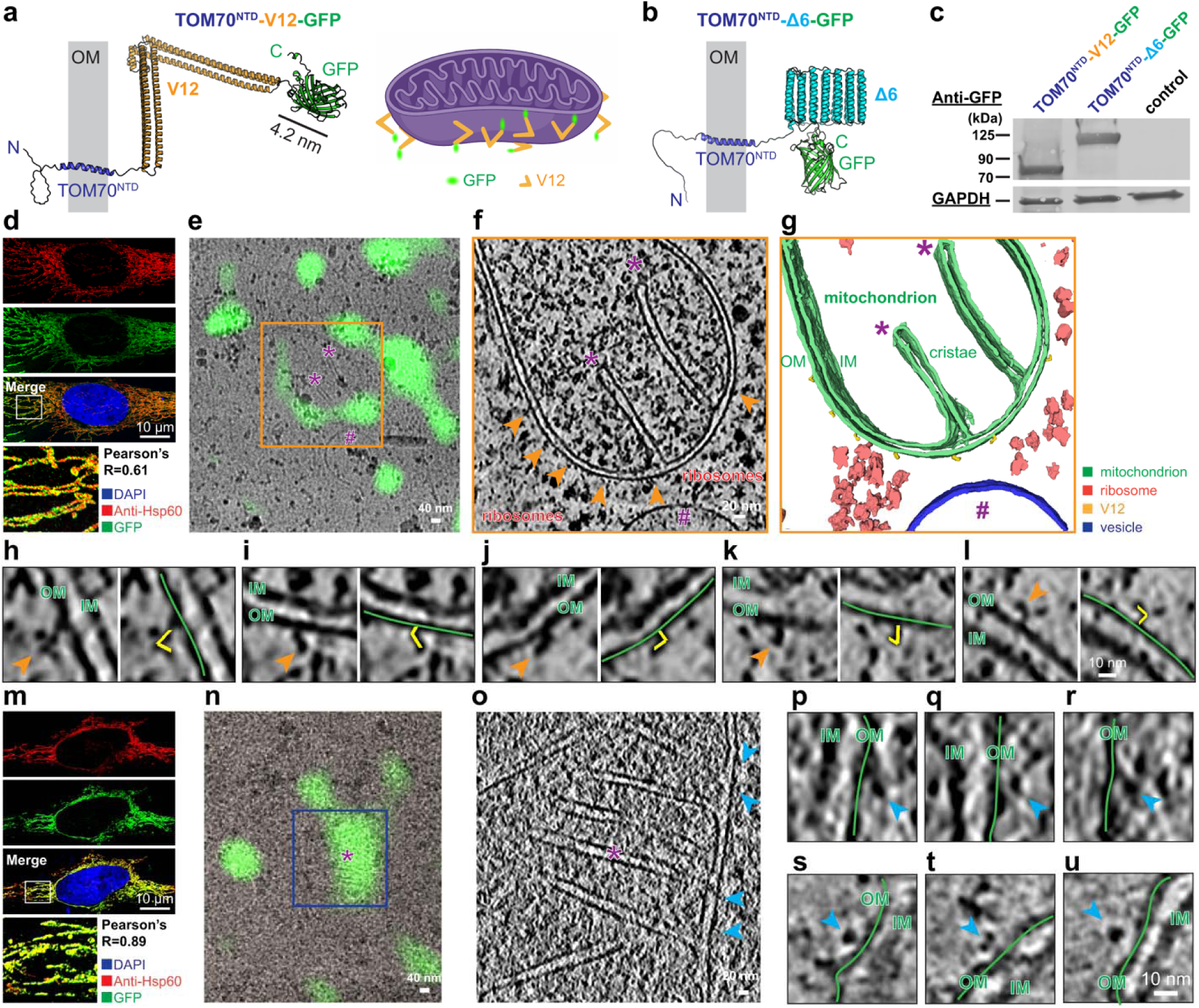
Mitochondrial surface display of V12- and **Δ**6-tagged TOM70^NTD^ fusion proteins in HeLa cells. (a, b) Schematic of TOM70^NTD^-V12-GFP and TOM70^NTD^-Δ6-GFP constructs. The TOM70 N-terminal domain (TOM70^NTD^) anchors to the mitochondrial outer membrane (OM), positioning the V12 or Δ6 tags on the cytosolic face. (c) Immunoblot of HeLa cell lysates expressing TOM70^NTD^-V12-GFP or TOM70^NTD^-Δ6-GFP probed with anti-GFP antibody. GAPDH served as a loading control. (d and m) Confocal fluorescence images showing mitochondrial localization of TOM70^NTD^-V12-GFP (d) and TOM70^NTD^-Δ6-GFP (m). GFP signal colocalizes with the mitochondrial marker Hsp60 (Pearson’s R = 0.61 and 0.89, respectively). Scale bars, 10 μm. (e) Cryo-correlative light and electron microscopy (cryo-CLEM) of TOM70^NTD^-V12-GFP cell. Fluorescence overlay shows GFP colocalized mitochondria on a FIB-milled lamella. (f) Tomographic slice of the corresponding region showing mitochondria, ribosomes, and cytosolic features; orange arrowheads indicate V-shaped densities. (g) Segmented tomogram showing mitochondria (green), ribosomes (red), and V12-tag densities (yellow). In panels e-g, purple asterisks (*) mark the same mitochondrial cristae, and purple hash symbols (#) mark the same vesicle. (h-l) Enlarged tomographic slices showing surface V12-tag densities (orange arrowheads) along the mitochondrial outer membrane (OM, green lines) and annotated V12 (yellow). IM, inner membrane. Scale bar, 10 nm. (n) Cryo-CLEM of TOM70^NTD^-Δ6-GFP cell. Fluorescence overlay shows GFP colocalized mitochondria on a FIB-milled lamella. (o) Tomographic slice of the corresponding region showing mitochondrion and cytosolic features; blue arrowheads indicate triangular-shaped densities. In panels N and O, purple asterisks (*) mark the same mitochondrion. (p-u) Enlarged tomographic slices showing compact Δ6-tag densities (blue arrowheads) on the mitochondrial outer membrane (green lines) of TOM70^NTD^-Δ6-GFP cells. OM, outer membrane; IM, inner membrane. Scale bar, 10 nm.

HeLa cells were transiently transfected; and GFP-positive cells were isolated by fluorescence-activated cell sorting (FACS), allowed to attach onto EM grids, and plunge-frozen for cryo-FIB milling (Extended Data Fig. 9). Cryo-fluorescence imaging of the resulting lamellae guided cryo-ET data acquisition and tracking of the tag (Fig. 4e, Extended Data Fig. 9). We reconstructed high-quality tomograms; after missing-wedge compensation and denoising with IsoNet^52^, V12-expressing cells showed well-resolved mitochondria, ribosomes, and vesicles (Fig. 4f, g). Cryo-fluorescence correlated with the 3D tomograms, revealing the signal on the mitochondrial surface (Fig. 4e). On the mitochondrial outer membrane, extended densities with the characteristic V-shaped geometry were clearly visible and annotatable, enabling precise 3D mapping of tag distribution (Fig. 4g, h-l and Supplementary Video 5).

In mito-Δ6-GFP expressing cells, ∼6 nm dot-like densities were observed on the mitochondrial surface and colocalized with fluorescence, with orientation-dependent appearances consistent with a compact triangular tag (Fig. 4n-u). However, these features were less distinct than those of V12 and difficult to assign unambiguously without reference. No V-shaped densities were detected in mito-Δ6 tomograms, further underscoring the uniquely identifiable morphology of the V12 tag.

Together, these results demonstrate that the V12 tags produce clear, detectable densities on mitochondrial surface in mammalian cells, correlates well with the GFP fluorescence signal. In contrast, the smaller and more compact Δ6 tag is challenging to resolve in situ without supporting experiments or subtomogram averaging results, consistent with the relative detectability observed in bacterial cells.

## Discussion

We introduce a fully genetically encoded, shape-defined tagging strategy based on a “shape-as-signal” principle, enabling direct identification of specific proteins in cryo-electron tomograms without post hoc labeling or chemical targeting. By encoding geometry rather than contrast, these low-molecular-weight, single-chain, monomeric tags form rigid, distinctive densities that are recognizable by eye at the electron microscope and amenable to computational validation. This creates a direct link between molecular identity and ultrastructural context—an essential step toward molecular-resolution maps of macromolecular organization in intact cells and toward routine in situ counting and positioning of individual proteins.

Compared with existing approaches such as nanogold labeling^21^, DNA-origami scaffolds^24^, or multimeric particles like GEMs^25^, our tags are fully genetically encoded, small enough to minimize perturbation of trafficking or localization, and engineered to fold into unambiguous 3D geometries. Their visibility arises not from contrast enhancement but from distinctive shape, analogous to how cytoskeletal filaments or membrane structures can be recognized in tomograms by morphology (size and shape) alone. Tagging ferritin or the mitochondrial outer membrane with either tag did not introduce detectable defects in protein assembly, trafficking, or morphology (Figs. 2-4). Notably, V12 is clearly visible on a standard 200-keV cryo-TEM (Glacios) in purified samples, and mammalian cells (Figs. 1, 4)—particularly in 3D tomograms—broadening accessibility and underscoring its potential for widespread application.

The two prototype designs illustrate a tunable design space. The extended V12 tag produces a characteristic V-shaped density that is readily detectable in situ on the mitochondrial outer membrane and in the cytoplasm of bacteria. The more compact Δ6 tag, although less visually striking in cells, is robustly identifiable in vitro. Together, these results suggest that tag geometry can be tailored to experimental needs—for example, maximizing detectability in crowded cytoplasm, minimizing footprint on a sensitive target protein, or introducing asymmetry so that the tagged terminus (N- or C-terminal) can be unambiguously assigned.

Beyond manual annotation, these tags have the potential to support automated analysis. In vitro, tagged complexes could be detected by both template matching^45,50^ and deep-learning based particle picking^43^, demonstrating feasibility for computational identification (Figs. 2, 3, and Extended Data Figs. 6, 7). Extending these approaches in situ should enable automated recognition of specific tagged molecules directly in cells. In particular, developing 3D (not merely 2D) detection algorithms specialized for V-shaped densities would improve recall and precision for low-molecular-weight features and accelerate both particle picking and subtomogram averaging, enabling automated detection and statistical analysis without requiring subtomogram averaging.

In cells, V12 could be directly recognized in tomograms and correlated with fluorescence signals from fusion to a fluorescence protein (e.g., GFP), allowing precise 3D mapping of its distribution on mitochondria. This ability to annotate the tagged protein’s position within its native ultrastructural environment creates a route to follow how localization changes across conditions such as signaling states, metabolic stress, or disease-associated mutations. More broadly, this bridges the LM-EM resolution gap: light microscopy provides temporal context and molecular specificity, while cryo-ET supplies molecular-resolution ultrastructure in the same cell with the exact same V12-FP fusion tag, without relying on CLEM post hoc physical correlation.

Looking forward, the protein-origami design framework is inherently extensible. Engineering additional tags with distinct, non-overlapping geometries would enable multiplexed labeling of different proteins in the same cell, allowing simultaneous mapping of multiple targets in 3D. In parallel, incorporation of heavy-atom clusters or tailored mass distributions could further improve detectability and support automated in situ identification.

This work is primarily a proof-of-concept demonstration of shape-defined, genetically encoded EM tags, and we do not yet use the approach to derive new biological insights. Our experiments establish feasibility in selected test systems, but each future application will require empirical optimization of tag placement, linker design, and expression levels, as well as functional controls to verify that the fusion does not perturb the behavior of the protein of interest—analogous to the validation routinely performed for fluorescent protein fusions. In addition, V-tags are currently identified mainly by visual inspection in cells and by simple template matching or deep-learning based particle picking in vitro. Nonetheless, our data show that V-shaped densities are readily identifiable in 3D volumes in situ, suggesting that robust automated detection is likely achievable. To fully realize large-scale, quantitative “visual proteomics,” dedicated 3D detection algorithms tailored to V-shaped densities will need to be developed and integrated into tomogram analysis pipelines.

In summary, these shape-specific, genetically encoded EM tags provide proof-of-principle for a general strategy to assign molecular identity directly in cryo-electron tomograms, practically bridging fluorescence imaging (temporal control, live-cell specificity, etc.) and cryo-ET (molecular-resolution ultrastructure), enabling direct tracking of protein localization under physiological and disease-relevant conditions. This approach opens a route to quantitative, context-aware maps of protein localization, organization, and interaction networks inside intact cells, laying the groundwork for truly integrative, in situ structural and functional proteomics.

## Materials & Correspondence

Supplementary Information is available for this paper.

Peer review information includes the names of reviewers who agree to be cited and is completed by Nature staff during proofing.

Reprints and permissions information is available at www.nature.com/reprints.

## Data and code availability

All cryo-EM/cryo-ET data will be deposited in EMPIAR (accession to be provided upon acceptance). The density maps and structure coordinates have been deposited in the EMDB and PDB under accession numbers EMD-73933 and 9Z9D (V12 tag) and EMD-73947 and 9Z9I (Δ6 tag). The original and/or analyzed data sets generated during the current study are available from the corresponding author upon reasonable request.

This paper does not report original code.

Any additional information required to reanalyze the data reported in this paper is available from the lead contact upon request.

## Acknowledgements

We are grateful to Drs. David Miller, Ege Kavalali, Lisa Monteggia, Borden Lacy, Hassane Mchaourab, Ian Macara (VU), Eric Skaar (VUMC), Z. Hong Zhou (UCLA) and Stella Sun (Pitt) for insightful discussions. We also thank Drs. Yun-Tao Liu and Hongcheng Fan (UCLA) for their support with IsoNet processing. EM data collection was performed at the Center for Structural Biology Cryo-EM Facility at Vanderbilt University. We acknowledge use of the Glacios cryo-TEM, which was acquired under NIH award S10 OD030292. Flow cytometry experiments were carried out in the VMC Flow Cytometry Shared Resource, which is supported by the Vanderbilt Ingram Cancer Center (P30 CA68485) and the Vanderbilt Digestive Disease Research Center (DK058404). Cryo-FIB milling was conducted at the Vanderbilt Institute of Nanoscale Science and Engineering with technical support from Dr. James McBride. Cryo-CLEM and cell imaging studies were performed in part through the Vanderbilt Cell Imaging Shared Resource, supported by NIH grants CA68485, DK20593, DK58404, DK59637, and EY08126. This work was supported by CDB Destination Postdoc Award to F.L., and grant from the National Institute of Health (R01MH132918 to Q.Z.).

## Author contributions

Conceptualization: F.L., Q.Z.; Methodology: F.L., R.S., O.C., P.L., Q.Z.; Investigation: F.L., R.S., P.L., Q.Z.; Visualization: F.L., Q.Z.; Funding acquisition: Q.Z.; Project administration: Q.Z.; Supervision: Q.Z.; Writing – original draft: F.L., Q.Z.; Writing – review & editing: F.L., R.S., O.C., P.L., Q.Z.

## Methods

### Protein Design and Computational Modeling of V- and **Δ**-shaped Tags

All shaped tags were designed as single-chain proteins with rigid, predefined geometries. We used an iterative, AlphaFold2-guided protein-engineering workflow (“protein nanoblocks”/Lego strategy): initial designs were modeled in AlphaFold2^40,54^, inspected for geometry and confidence, and refined through successive design–prediction cycles (Extended Data Fig. 3). All surface residues were tuned for hydrophilicity. Electrostatic surface potentials were calculated in PyMOL (APBS plugin) ^55^ to verify balanced charge distribution across exposed surfaces and to reduce the risk of nonspecific interactions or oligomerization.

For the V-shaped protein, the V scaffold was derived from a three-helix-bundle (PDB: 4TQL) with the two bundles connected by a rigid turn inspired by sterile α-motif (SAM) domains^38,56,57^ and a *de novo*-designed mini-protein motif^58^. AlphaFold2 predicted four candidates with inter-arm angles of ∼60°, 72°, 90°, and 140° (Extended Data Fig. 1). To maintain solubility and prevent oligomerization or undesired interactions, SAM-interface residues were neutralized.

For the Δ-shaped protein, we used the same design strategy, Δ6 was built from a C3-symmetric trimeric scaffold (*C3*triangle120_C3_A) to form an equilateral triangular assembly (∼6 nm per side)^41^. Two short linkers were engineered to concatenate three repeats into a single chain, preserving the C3 geometry.

### Protein Expression and Purification

For V12 and Δ6 proteins, codon-optimized genes encoding V12 and Δ6 were cloned into pET27b vectors with N-terminal His_6_ tags for expression in *E. coli* BL21(DE3) (NEB). Cultures were grown in LB at 37°C to OD_600_ ≈ 0.6, induced with 0.1 mM isopropyl-β-D-thiogalactoside (IPTG), and incubated for 12 h at 20°C. Cells were pelleted and resuspended in lysis buffer (20 mM Tris-HCl pH 8.0, 300 mM NaCl, 10 mM imidazole) supplemented with a protease inhibitor cocktail tablet (Roche). After sonication and centrifugation (18,000 × g, 60 min) at 4°C, supernatants were purified by Ni–NTA affinity chromatography (Ni-NTA Agarose, Qiagen), anion-exchange chromatography (Resource Q, Cytiva), and size-exclusion chromatography (Superdex 200 Increase 10/300 GL, Cytiva) in 20 mM Tris-HCl pH 8.0, 300 mM NaCl. Protein fractions were verified by SDS-PAGE and concentrated to ∼0.5 mg/mL for cryo-EM.

For V12-ferritin and Δ6-ferritin nanocages, the *E. coli* ferritin (*ftnA*) gene was fused at its N terminus to either V12 or Δ6 via a flexible linker and were cloned into pJ414 vectors with N-terminal His_6_ tags. Cultures were grown and induced with as above but harvested after 4 h at 20°C. A portion of each culture (1 mL) was used directly for plunging freezing and cryo-FIB milling. The remaining cells were pelleted, resuspended in lysis buffer (20 mM Tris-HCl pH 7.4, 300 mM NaCl, 10 mM imidazole), supplemented with a protease inhibitor cocktail tablet (Roche) at 4°C. The cells were lysed by sonication, and clarified by centrifugation (18,000 × g, 60 min) at 4°C. Purification followed the same chromatography workflow as above with the buffer at pH 7.4. Purified samples were verified by SDS-PAGE and concentrated to ∼0.5 mg/mL for cryo-EM.

### Single particle cryo-electron microscopy

Purified V12 and Δ6 proteins were applied to glow-discharged Quantifoil R1.2/1.3 Cu 300-mesh grids and vitrified using a Vitrobot Mark III (FEI) (95% humidity, 4°C, blot time 3 s). Data were acquired on a 200-keV Thermo Fisher Glacios TEM equipped with a Falcon 4 direct detector at 120,000× magnification (pixel size 0.73 Å) with a total dose of 60 e⁻/Å^2^ in EER format. Beam induced motion-correction and dose-weighting to compensate for radiation damage over spatial frequencies were performed using Patch Motion correction and Contrast Transfer Function (CTF) estimation were performed in cryoSPARC^59^. Particle picking, two-dimensional (2D) classification, and 3D refinement produced final reconstructions, reached overall resolutions of 5.7 Å for V12 and 6.8 Å for Δ6 by gold-standard Fourier shell correlation (FSC) at the 0.143 criterion. Both datasets were processed without applying symmetry (C1), allowing unbiased reconstruction of the full asymmetric architectures of the tags.

### Mammalian cell culture, transfection, and FASC

HeLa (ATCC, no. CCL-2) and HEK293T (ATCC, no. CRL-3216) were cultured in DMEM (Gibco, no. 31053028) supplemented with 10% (v/v) fetal bovine serum (FBS, Gibco, no. A5669701), and 1% MEM nonessential amino acids (Gibco, no. 11140-050) at 37°C with 5% CO_2_.

For mitochondrial targeting, TOM70^NTD^-V12 and TOM70^NTD^-Δ6 constructs tagged with GFP or HA were cloned into pFUGW backbone under the UBC promoter. TOM70^NTD^ corresponds to residues 1-59 of human TOM70 protein, which mediates outer mitochondrial membrane localization.

Cells were seeded into 10 cm dishes one day before transfection. At ∼70% confluency, transfections were performed using FuGENE 6 (Promega, no. F6-1000) with 5µg of plasmids DNA and Opti-MEM (Gibco, no. 31985062) following the manufacturer’s protocol. Two days post-transfection, GFP-positive cells were sorted by flow cytometry using a BD FACS Aria III. Parallel transfections were carried out in 6-well or 24-well plates for immunoblotting and immunofluorescence assays.

### Cryo-ET sample preparation

For *E. coli* expressing V12-ferritin and Δ6-ferritin, *E. coli* cultures (1mL) expressing V12-ferritin or Δ6-ferritin (described above) were centrifuged at 2500 × g for 5min, washed once with PBS (pH 7.4) and resuspended into ∼60 µL PBS. Cell suspensions were applied to glow-discharged Quantifoil R2/2 Cu 200-mesh grids and plunge-frozen using a Vitrobot Mark III (FEI) at 95% humidity and 24°C with a 3s blot time.

For HeLa cell preparation, Gold Quantifoil R2/2 SiO_2_ film grids were UV-sterilized for 30min per side and coated with sterilized 0.05 mg/mL poly-L-lysine (PLL, Sigma-Aldrich, no. P2636-100MG) in 0.1M borate buffer (pH 8.5; Boric Acid, Sigma-Aldrich, no. B-0252; Borax, Sigma-Aldrich, no. B-9876) overnight at room temperature. Grids were rinsed 3 times with ddH_2_O and equilibrated in culture medium.

After cell sorting, GFP-positive cells were pelleted with 200 × g for 5 min and resuspended in medium containing 4 µM AraC (to prevent division) and HEPES and seeded onto 3-well dishes (Culture-Insert 3 Well in 35 mm µ-Dish, ibidi, no. 80366) with the amount of ∼1× 10^4^ cells per 70 µL with 2 grids each well. Six hours after attaching, grids cultured with GFP-positive HeLa cells were plunge-frozen in pre-warmed PBS using Leica EM GP2 with one side blotting at 37°C, 95% humidity, 3 s blotting time.

### Immunoblotting and immunofluorescence

Cells were lysed in RIPA buffer (25 mM Tris pH 7.6, 150 mM NaCl, 1% NP-40; Sigma, no. R0278) supplemented with protease inhibitors. Lysates were separated by SDS-PAGE using 4%-20% Mini-PROTEIN TGX Precast Protein Gels (Bio-RAD, no. 4561094) and transferred to PVDF membranes. Immunoblotted was performed with anti-GFP (Roche, no. 11814460001, 1:1,000) or anti-HA (Invitrogen, no. 26183, 1:5,000) primary antibodies, and GAPDH (Cell signaling, no. 2118S, 1: 1,000) served as a loading control. IRDye secondary antibodies (LI-DOR) were used for detection, and signals were imaged with an Odyssey DLx system (LI-COR).

For immunofluorescence, cells were fixed with 4% paraformaldehyde (PFA), permeabilized with 0.1% Triton X-100, and stained with anti-HA (Invitrogen, no. 26183, 1:500; magenta), anti-Hsp60 (Cell signaling, no. 12165S, 1:200), MitoTracker Red CMXRos (Invitrogen, no. M46752), and DAPI (blue). Images were acquired using a Nikon CSU-W1 SoRa confocal microscope and Nikon SIM system. Colocalization with mitochondria was quantified in FIJI^60^ using Pearson’s correlation coefficient.

### Cryo-FIB lamella preparation

Cryo-focused ion beam (cryo-FIB) milling was performed using an FEI Helios NanoLab G3 CX with a Quorum PP3010T cryo-SEM system at liquid nitrogen temperature. Prior to milling, metallic platinum was deposited by sputter coating (10 mA, 20 s), followed by a protective layer of organometallic platinum applied via the gas injection system (6 mm working distance, 25° stage tilting angle and 8s injection).

Two notches were first created ∼1 μm away from the lamella to relieve mechanical stress and prevent warping or bending during subsequent thinning and transfer. Cells were then milled to ∼1 μm thickness at a 20° stage tilt using ion beam currents of 0.43 nA and 0.23 nA at 30 keV. The stage was then tilted to 16°, and lamellae were thinned to a target thickness of 400–500 nm using beam currents of 80 pA and 40 pA. Finally polishing was performed at 16° with cross-cleaning at 23 pA to achieve a final thickness of 100-250 nm. Before unloading, SEM overview image of all lamellae and the corresponding grid was acquired to provide localization references for subsequent cryo-CLEM. Finally, lamellae were sputter-coated with platinum (3 mA, 2 s) to minimize charging and beam-induced drift during cryo-ET imaging.

### Cryo-correlative light and electron microscopy (Cryo-CLEM)

Cryo-FIB-milled lamellae of HeLa cells expressing TOM70^NTD^-V12-GFP or TOM70^NTD^-Δ6-GFP were imaged using Leica STELLARIS Cryo-confocal microscope. FIB-milled grids were transferred with a Leica EM VCM under fresh liquid nitrogen to limit ice containment.

Lamellae were first located in widefield mode based on overview SEM reference images. Subsequently, z-stacks encompassing the entire lamellae and adjacent notches were acquired in Lighting mode using 491 nm and 587 nm lasers to capture GFP fluorescence and autofluorescence, respectively, for later correlation with TEM search maps. Z-stacks were processed to generate sum-intensity projections. Correlation between cryo-fluorescence images and low-magnification TEM search maps (lamella overviews) was performed using IMOD^61,62^ and FIJI^60^.

### Cryo-ET image acquisition

For purified V12-ferritin and Δ6-ferritin nanocages, the purified samples were applied to glow-discharged Quantifoil R2/2 Cu 200 mesh grids and plunge-frozen as described above. Tilt series were collected from −55° to +55° in 5° increments with dose-symmetric tilt scheme (8e^-^/Å^2^ per tilt; total accumulated dose ∼184 e^-^/Å^2^) on a 300 kV Titan Krios G4 microscope equipped with a Gatan K3 detector and a BioQuantum energy filter. Data were acquired at a nominal defocus of 3-4 µm, using Thermo Fisher Tomography software.

For bacterial and mammalian lamellae, the stage was tilted by ±9° to compensate for the final milling angle. Tilt series were collected from −60° to +60° using a dose-symmetric tilt scheme with 2° increments (total dose ∼183 e⁻/Å^2^). The *E. coli* lamellae were imaged on a 300 kV Titan Krios G4 microscope equipped with a Gatan K3 detector and energy filter, using a defocus of 3-5 µm and a calibrated pixel size of 1.6 Å. HeLa cell lamellae were first screened by collecting low-magnification search maps for all existing lamellae. Cryo-fluorescence correlation with CLEM data was performed as described above to identify regions containing both GFP signal and mitochondria for targeted cryo-ET data acquisition. Tilt series were collected on a 200 kV Thermo Fisher Glacios TEM equipped with a Falcon 4 direct detector, using 4-5 µm defocus, a 70 µm objective aperture, and a pixel size of 1.5 Å.

### Cryo-ET data processing

For purified V12-ferritin and Δ6-ferritin nanocages, tilt series were aligned and reconstructed in RELION5^47,63^ with integrated motion correction and CTF correction. Reconstructed tomograms were binned fourfold and processed with IsoNet for missing-wedge compensation and denoising, enabling improved model fitting and visualization.

Subtomogram averaging (STA) was performed using crYOLO^43,44^ for automatic ferritin cage picking and PyTom^45^ for localization of smaller tag particles. Amond tested approaches, crYOLO^43,44^ was most effective for large in vitro particles, whereas PyTom^45^ performed better for small tag features in vitro and in situ cage detection. Independent refinements of cage and tag subtomograms were carried out in RLION5^47,64,65^, yielding final resolution of 5 Å and 22 Å for ferritin cage and the V12 tag, respectively, and 6.7 Å and 7.3 Å for ferritin cage and Δ6 tag, respectively. Averaged densities were fitted into corresponding tomograms using UCSF ChimeraX^66^ for visualization, tags detection and structural interpretation.

For Bacterial and mammalian cell tomograms, tilt series of *E. coli* and HeLa cell lamellae were motion corrected with Motioncor3^67^ and reconstructed using IMOD (weighted back-projection mode)^61,62^ and binned fourfold, yielding final pixel size of 6.4 Å (*E. coli*) and 6 Å (HeLa). The tomograms were subsequently processed with IsoNet^52^ for missing-wedge compensation and denoising, using custom masks generated to focus on regions enriched in ferritin cages or mitochondrial membranes and associated tags. Ribosomes, membranes, and ferritin nanocages were segmented using AI-assisted tools in Amira (Thermo Fisher Scientific).

Tag-like densities were identified through manual inspection and validated by docking averaged tag models obtained from purified samples into tomographic volumes using ChimeraX^66^. While PyTom^45^ enabled efficient in situ cage picking, existing algorithms failed to reliably detect the smaller tag densities due to the combination of the missing wedge and the crowded cellular environment. STA of in situ ferritin cages, performed using Warp^46^ and RELION5^63,64^, achieved a final resolution of ∼12 Å.

Current algorithmic limitations hinder robust automated identification of small, shape-defined tags in situ. Ongoing efforts aim to develop new computational approaches tailored for these geometrically defined tags to enhance their detection and verification within cellular tomograms. Although technically challenging, such advancements are expected to substantially broaden the applicability and usability of both tags in future studies.

### Data analysis and visualization

All density maps were visualized in UCSF ChimeraX^66^ and segmented in Amira (Thermo Fisher Scientific). Electrostatic potential surfaces were rendered in PyMOL with APBS^55^. Fourier shell correlation (FSC) was used to estimate resolution^68^. For 3D modeling, structures were fitted into tomograms using ChimeraX^66^. Figures were prepared in ChimeraX, PyMOL, BioRender (Extended Data Figs. 3, 9A), and Adobe Illustrator.

