## Supplemental information for "Fully genetically encoded low-molecular-weight protein tags with defined shapes for direct molecular identification by cryo-electron tomography"

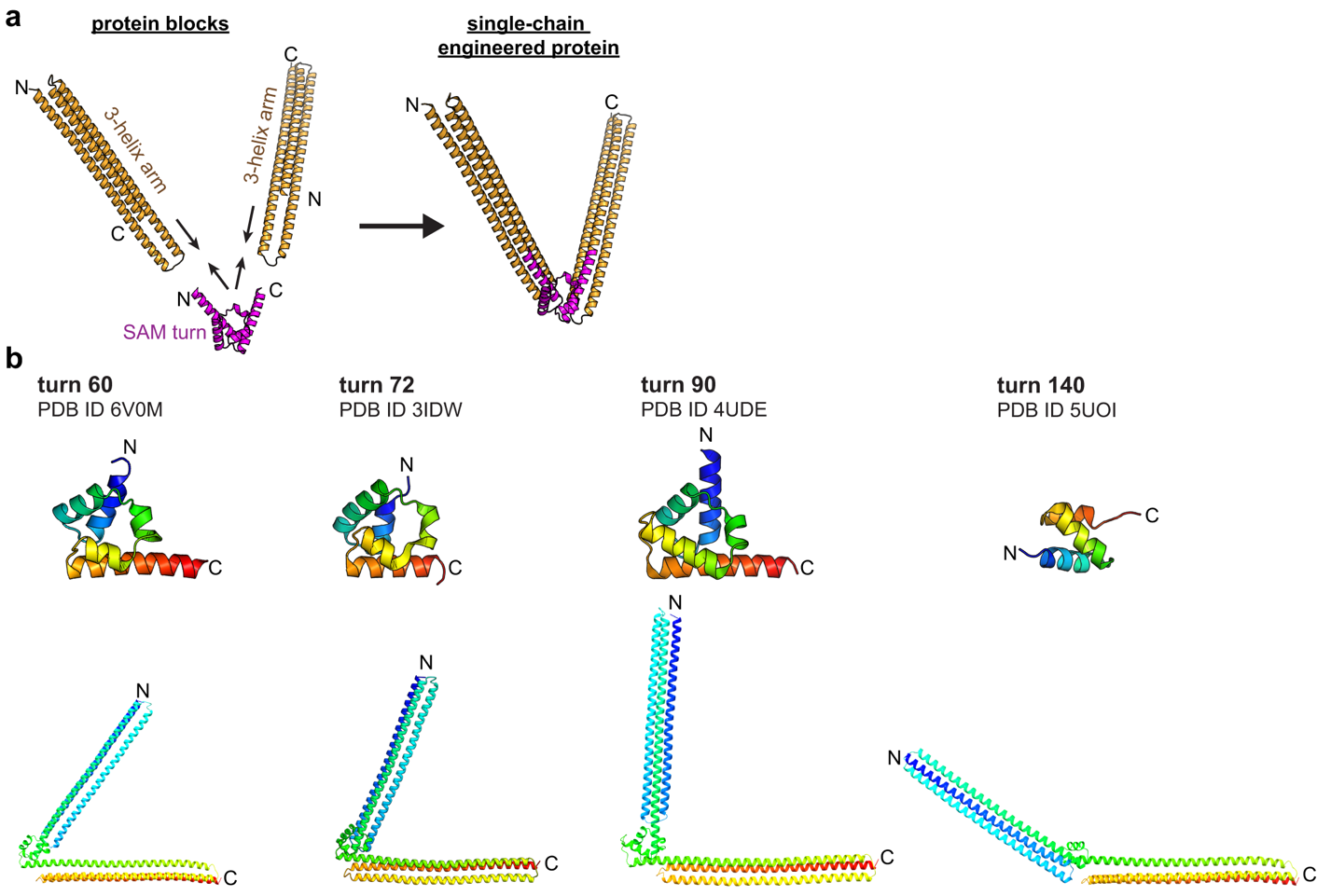

Extended Data Fig. 1. Design and computational validation of the V-shaped protein tag.

(a) Schematic showing the modular design strategy. Two three-helix bundle arms were connected through a rigid SAM-turn motif to form a single-chain, V-shaped protein with a defined inter-arm angle.

(b) Predicted models of four V-shaped candidates with different inter-arm angles (~60°, 72°, 90°, and 140°) based on distinct SAM-turn motifs (PDB IDs: 6V0M, 3IDW, 4UDE) and a *de novo*-designed mini-protein motif (PDB ID: 5UOI) (from David Baker’s lab). The top panels show the corresponding SAM-turn structures; the bottom panels show the full V-shaped designs predicted by AlphaFold2.

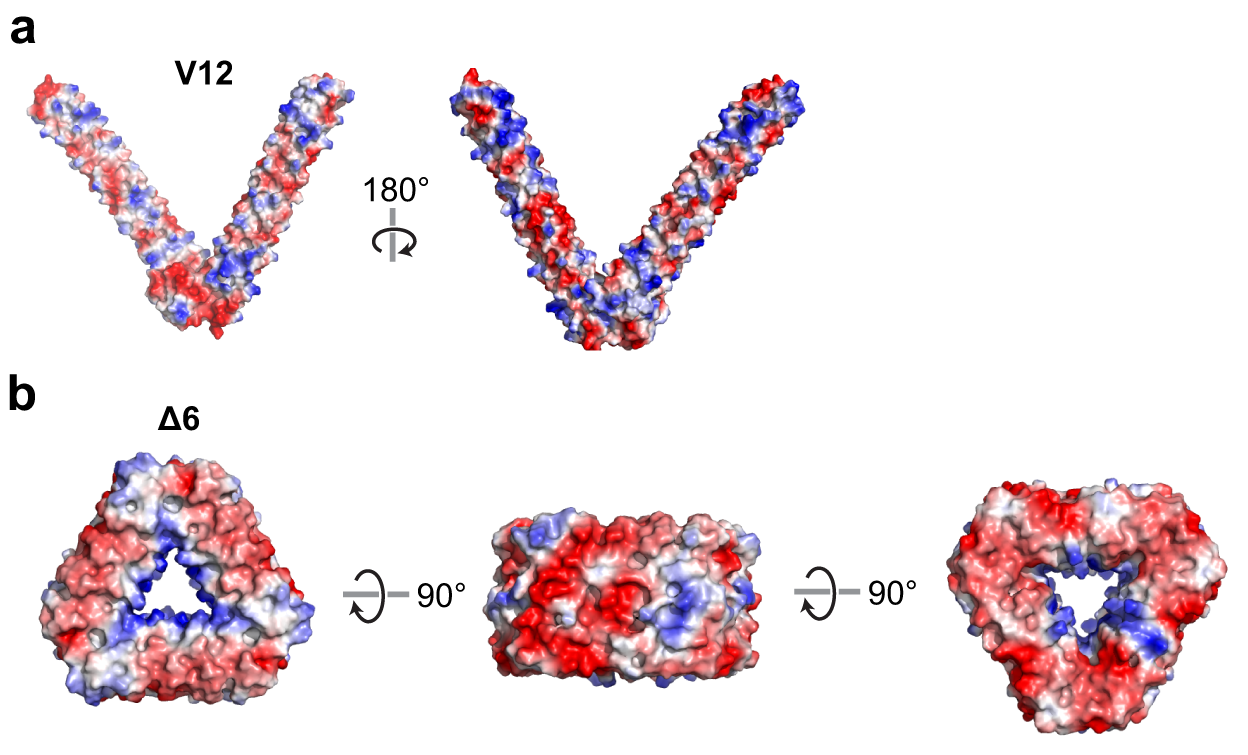

Extended Data Fig. 2. Electrostatic surface potential of V12 and Δ6 tags.

(a) Electrostatic surface map of the V12 protein viewed from two orientations (rotated 180°). The surface shows alternating positively (blue) and negatively (red) charged regions distributed along the two helical arms.

(b) Electrostatic surface maps of the Δ6 tag viewed from three orientations (rotated 90°). The trimeric assembly displays a balanced charge distribution across the compact triangular surface and central pore, consistent with its stable oligomeric architecture.

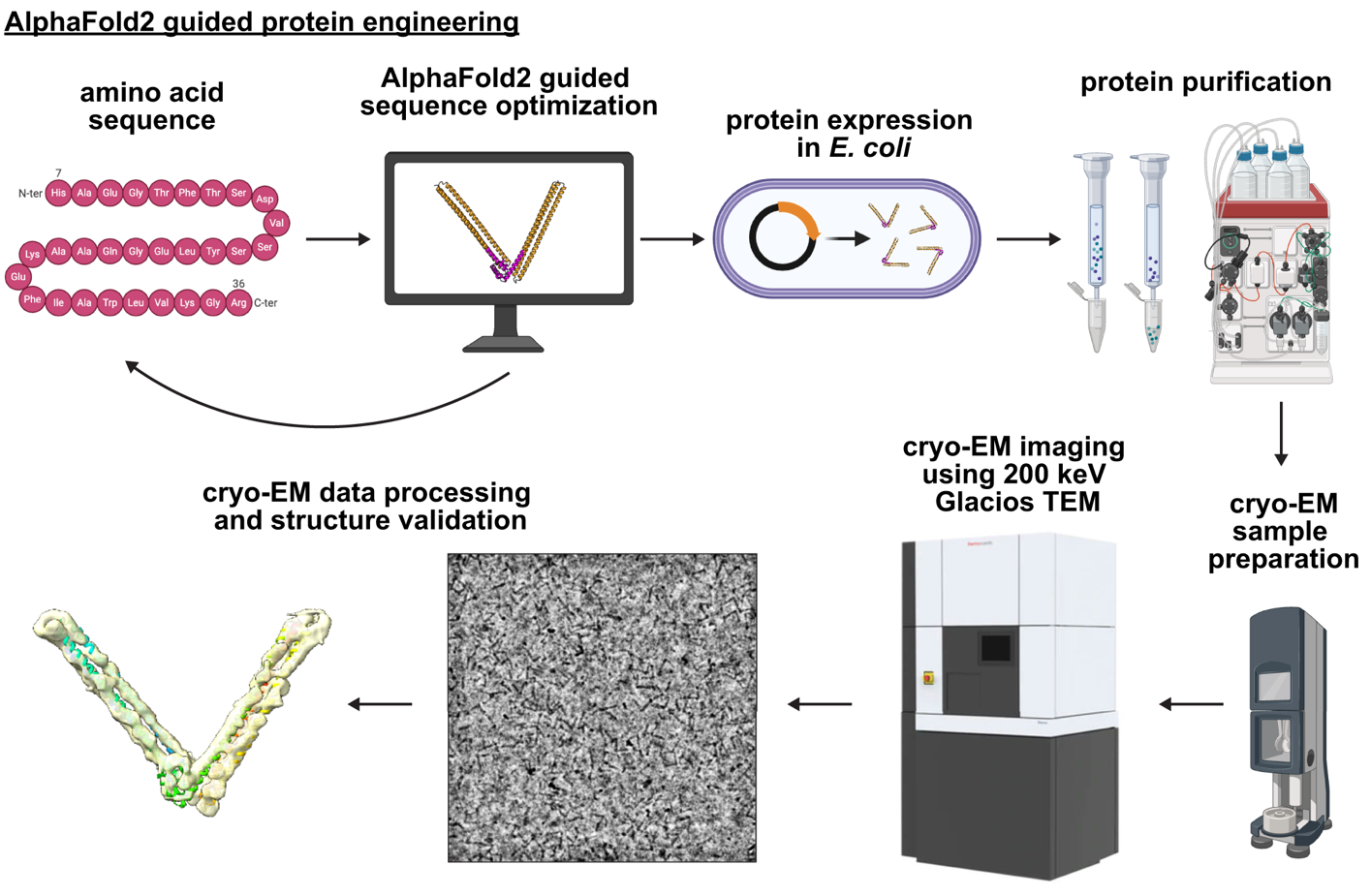

Extended Data Fig. 3. Workflow of AlphaFold2-guided protein engineering.

Predicted models were refined through sequence optimization, followed by expression in *E. coli*, purification, cryo-EM imaging using a 200-keV Glacios TEM, and structural validation by cryo-EM reconstruction.

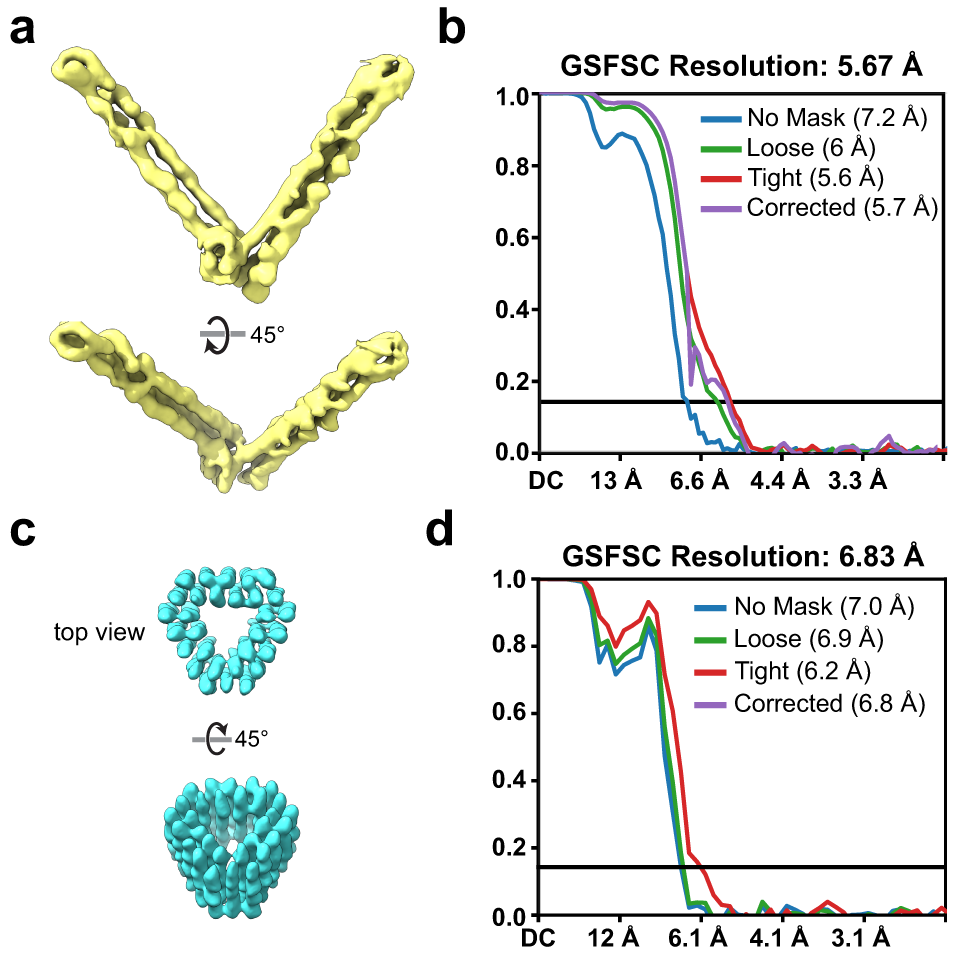

Extended Data Fig. 4. Cryo-EM density maps and resolution estimation of purified V12 and Δ6 tags.

(a) Cryo-EM density map of the extended V12 tag viewed from two orientations (rotated 45°).

(b) Gold-standard Fourier shell correlation (GSFSC) curves showing estimated resolutions for V12 under different masking conditions, with the final resolution of 5.67 Å.

(c) Cryo-EM density map of the compact Δ6 tag viewed from top and 45°-rotated orientations. (d) GSFSC curves showing estimated resolutions for Δ6 under different masking conditions with a final resolution of 6.83 Å.

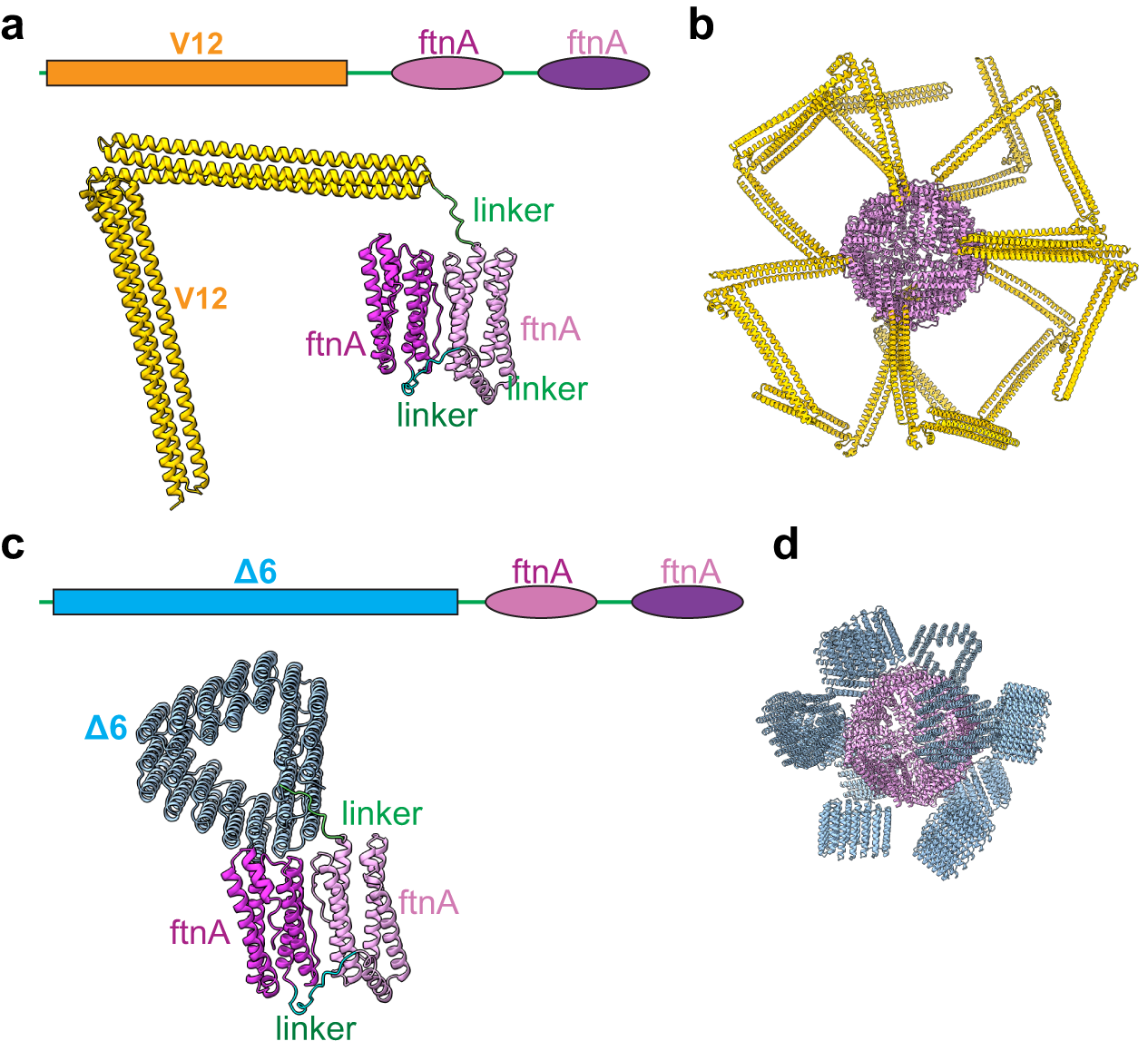

Extended Data Fig. 5. Structural models of V12- and Δ6-tagged ferritin nanocages.

(a) Schematic and structural model of the V12-ferritin (FtnA) fusion. Each ferritin subunit (pink) is connected to the V12 tag (yellow) through a flexible linker (green).

(b) Modeled assembly of the 24-meric ferritin cage decorated with extended V12 tags radiating outward.

(c) Schematic and structural model of the Δ6-ferritin fusion. Each ferritin subunit (pink) is linked to the compact Δ6 tag (blue) via a flexible linker (green).

(d) Modeled assembly of the 24-meric ferritin cage showing peripheral triangular Δ6 tags.

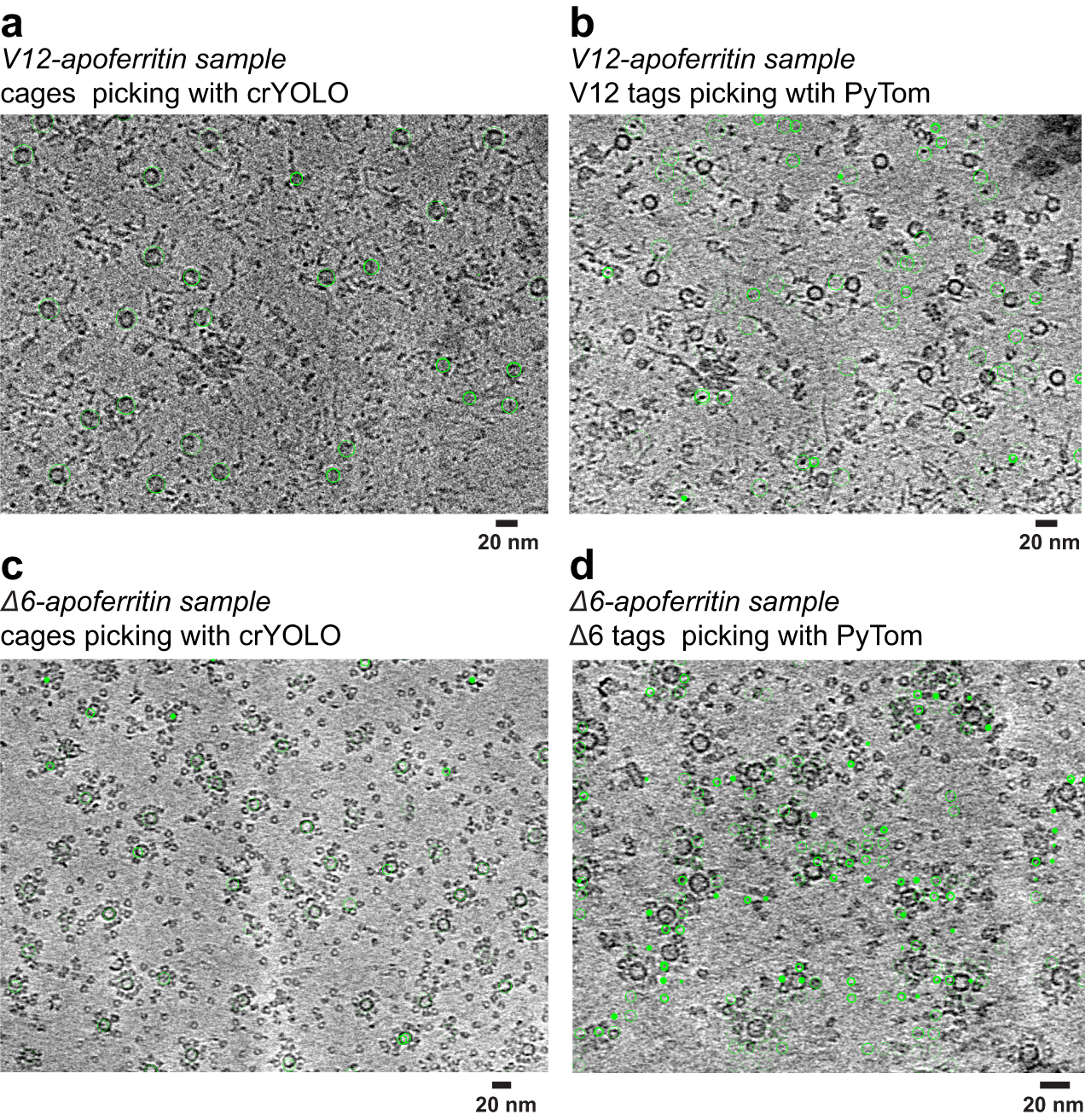
 **Extended Data Fig. 6.** **Particle picking of purified V12- and Δ6-tagged apoferritin samples from cryo-tomograms.**
(a, b) Representative tomographic slices of purified V12-apoferritin samples. (a) Ferritin cages were automatically picked using a deep-learning based program of crYOLO. (b) V12 tag densities were independently picked using template matching and correlation-based approach of PyTom. Green circles indicate selected particles.
(c, d) Representative tomographic slices of purified Δ6-apoferritin samples. (c) Ferritin cages were picked with crYOLO. (d) Δ6 tag densities were independently picked with PyTom. Green circles indicate selected particles. Scale bars, 20 nm.

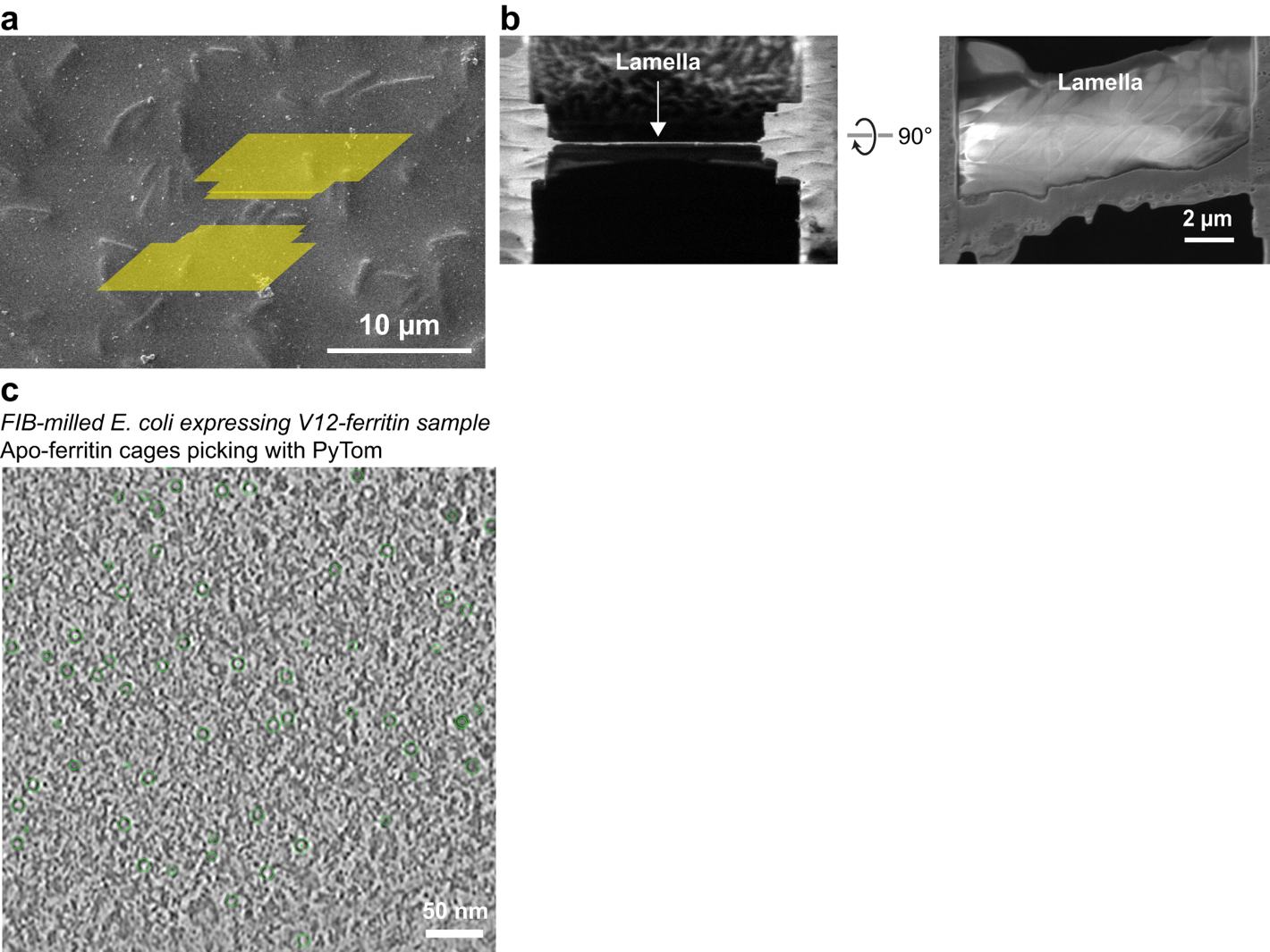

Extended Data Fig. 7. Cryo-FIB milling and tomogram-based particle picking of *E. coli* expressing V12-ferritin.

(a) SEM image showing the removed area (highlighted in yellow) prepared by cryo-focused ion beam (cryo-FIB) milling to generate thin lamellae of *E. coli* cells.

(b) Cross-sectional and side-view SEM images of a milled lamella. The lamella thickness was ~150 nm.

(c) Representative tomographic slice from the FIB-milled *E. coli* sample expressing V12-ferritin. Apoferritin cages were automatically picked using PyTom (green circles).

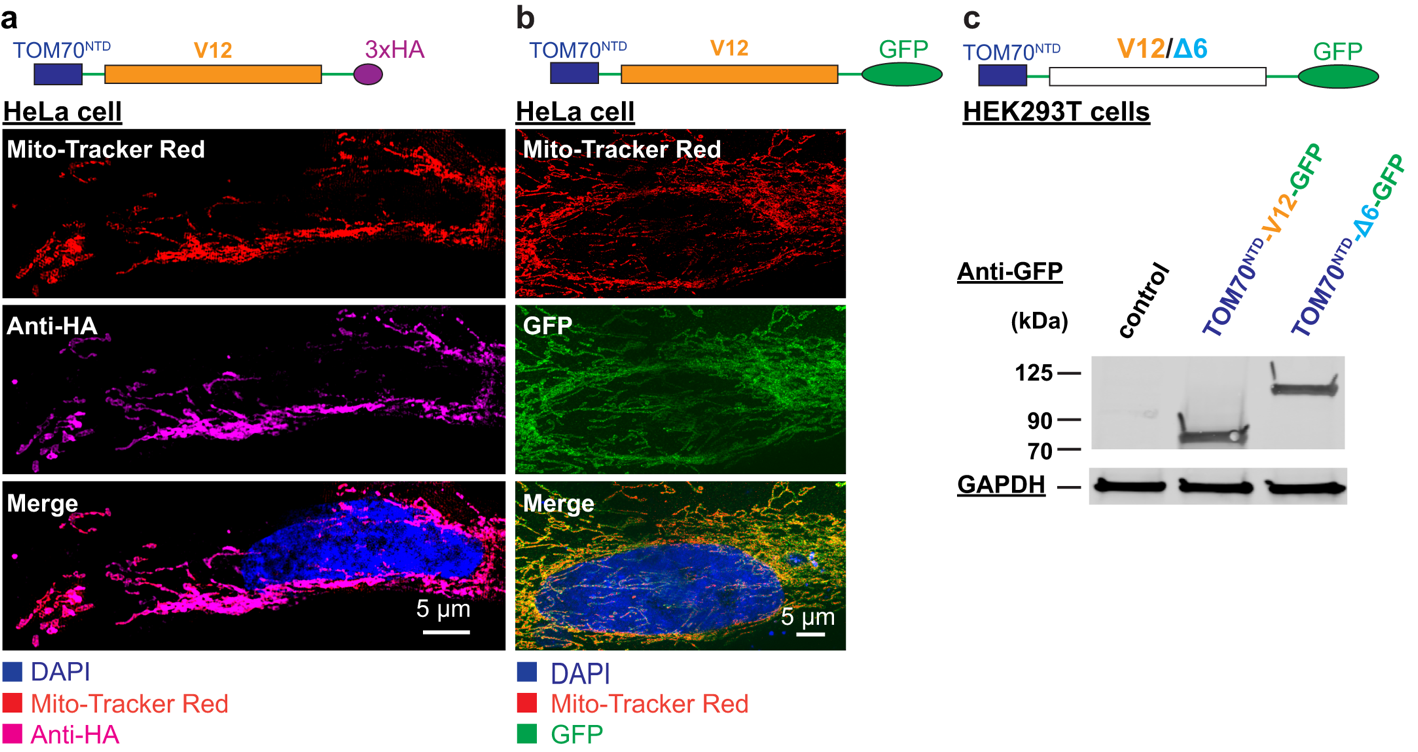

Extended Data Fig. 8. Expression and mitochondrial localization of TOM70^NTD^-V12/Δ6 constructs in mammalian cells.

(a) Schematic and fluorescence images of HeLa cells expressing TOM70^NTD^-V12-3×HA. Mito-Tracker Red (mitochondria) and anti-HA staining (magenta) show colocalization of the V12 fusion protein with mitochondria. Nuclei are stained with DAPI (blue).

(b) Schematic and fluorescence images of HeLa cells expressing TOM70^NTD^-V12-GFP. Mito-Tracker Red and GFP signals colocalize along mitochondrial networks, confirming proper targeting.

(c) Schematic of TOM70^NTD^-V12/Δ6-GFP constructs and immunoblot of HEK293T cell lysates probed with anti-GFP antibody. GAPDH serves as a loading control.

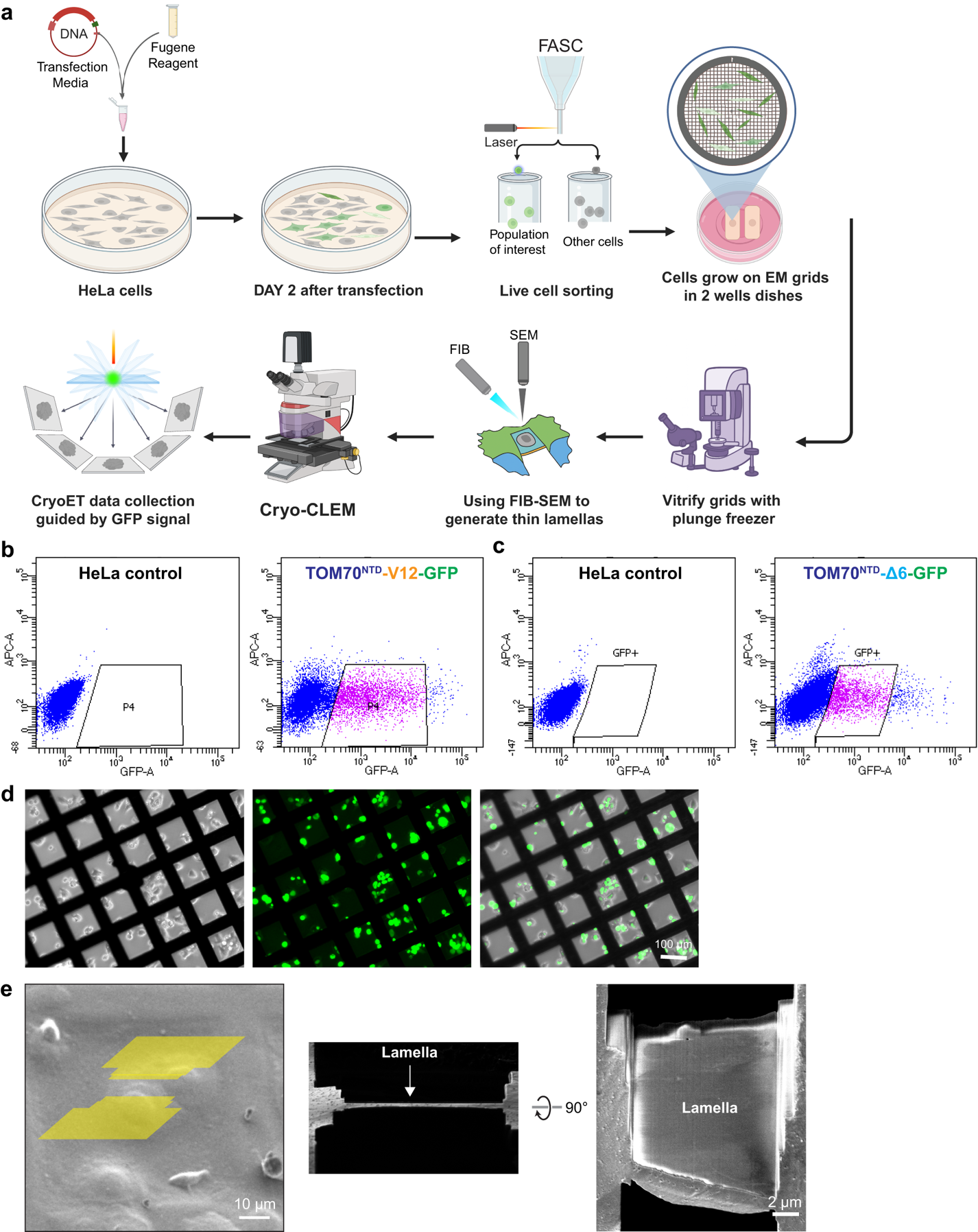
**Extended Data Fig. 9. Cryo-CLEM and cryo-FIB-SEM workflow for imaging TOM70^NTD^-V12/Δ6-GFP-expressing HeLa cells.**

(a) Schematic of the experimental workflow. HeLa cells were transfected with TOM70^NTD^-V12-GFP or TOM70^NTD^-Δ6-GFP constructs using FuGENE reagent. Two days post-transfection, cells were sorted by fluorescence-activated cell sorting (FACS) to enrich GFP-positive populations and then cultured and grown on EM grids in 2-well dishes. After cells were attached to EM grids, vitrified by plunge-freezing, and thin lamellae were prepared by cryo-focused ion beam-scanning electron microscopy (cryo-FIB-SEM). Cryo-correlative light and electron microscopy (cryo-CLEM) guided cryo-ET data collection.

(b, c) Flow cytometry plots of HeLa control and transfected cells showing GFP-positive populations for TOM70^NTD^-V12-GFP (b) and TOM70^NTD^-Δ6-GFP (c).

(d) Fluorescence images of EM grids before plunge-freezing showing uniform GFP-positive cell distribution and attachment.

(e) SEM images of cryo-FIB-milled lamellae of HeLa cells expressing TOM70^NTD^-V12-GFP or TOM70^NTD^-Δ6-GFP. Yellow overlays mark milled positions; cross-sectional and side-view images show a thin lamella suitable for tomography.

Supplementary Video 1. Fitted model of purified V12-ferritin nanocage

The model was generated by fitting the ferritin cage and V12 tag obtained from sub-tomogram averaging (STA) into the purified 3D tomographic density. Slice views demonstrate a good agreement between the model and the original density.

Supplementary Video 2. Fitted model of purified Δ6-ferritin nanocage

The model was generated by fitting the ferritin cage and Δ6 tag obtained from STA into the purified 3D tomographic density. Slice views demonstrate a good agreement between the model and the original density.

Supplementary Video 3. Fitted model of *E. coli* V12-ferritin nanocage

The model was generated by fitting the structures of in vitro ferritin cage and V12 tag obtained from STA into the in situ 3D tomographic density. Slice views demonstrate a good agreement between the model and the original density.

Supplementary Video 4. Fitted model of *E. coli* Δ6-ferritin nanocage

The model was generated by fitting the structures of in vitro ferritin cage and Δ6 tag obtained from STA into the in situ 3D tomographic density. Slice views demonstrate a good agreement between the model and the original density.

Supplementary Video 5. Tomogram and annotation of mitochondria within a HeLa cell expressing TOM70^NTD^-V12-GFP

The video presents the full tomogram of mitochondria, showing mitochondrial membranes, cristae, ribosomes, and surrounding cytosolic structures, with annotations highlighting prominent features and the distribution of V12 tags.
